# Ulinastatin prevents hyperoxia-induced lung injury on newborn rats by downregulating TNF-*α* and inhibiting macrophage infiltration

**DOI:** 10.1101/494773

**Authors:** Feng-dan Xu, Wen-long Deng, Xiang-yong Kong, Zhi-chun Feng

**Affiliations:** Department of Newborn Care Center, Affiliated BaYi Children’s Hospital, Clinical Medical College in PLA Army General Hospital, Southern Medical University, Beijing 100700, China; Department of Pediatrics, GuangDong Medical University, Dongguan 523808, Guangdong, China; Department of Critical Care Medicine, The third People’s Hospital of Dongguan City, Dongguan 523326, Guangdong, China

**Keywords:** Ulinastatin, hyperoxia-induced lung injury, bronchopulmonary dysplasia, anti-inflammation

## Abstract

Ulinastatin (UTI) can support protection for several organs through inhibition of the release of inflammatory factors, absorbing oxygen radicals, and inhibition the progression of fibrosis. However, whether UTI has effect on hyperoxia-induced lung injury is still unclear. In this study, the effects of UTI treatment on newborn rats suffering hyperoxia-induced lung injury were examined. The results demonstrated that UTI treatment significantly attenuated the wet/dry weight ratio, downregulated the levels of tumor necrosis factor-α (TNF-α), inhibited macrophage infiltration, and improved the average weight of rats. The most significant changes were observed in high-dose UTI treatment group. However, UTI had no effect on pulmonary superoxide dismutase (SOD) and malondialdehyde (MDA) levels. Our results suggested that Ulinastatin prevents hyperoxia-induced lung injury on newborn rats by downregulating TNF-α and inhibiting macrophage infiltration.

## Introduction

Approximately 30% of premature infants with birth weights <1,000 g develop bronchopulmonary dysplasia (BPD), which increases their morbidity and mortality and influences their long-term outcome.^1^ Although the pathogenesis of BPD is not completely understood, hyperoxia, inflammation, and/or ventilator exposure of premature lung are thought to be principal causative factors.^2^ More importantly, there is no effective clinical treatment for BPD.^3^

Ulinastatin, also known as urinary trypsin inhibitor (UTI), was first identified in and purified from human urine by a group of Japanese researchers in 1982.^4^ It has been reported that UTI have potent antioxidant, anti-inflammatory, and antimicrobial properties. UTI suppresses excessive radical superoxide anion generation in blood, oxidative stress in the forebrain of rats with ischemia/reperfusion.^5^ The enhanced production of pro-inflammatory mediators, including interleukin-8 (IL-8) and tumor necrosis factor-α (TNF-α), could be inhibited by UTI.^6,7^ Furthermore, it has been shown to inhibit the development of radiation-induced lung fibrosis in mice.^8^ However, it is not known if UTI has an effect on the development of BPD.

The current study focuses on the impact of UTI treatment on a rat model of hyperoxia-induced lung injury, which is a well-established experimental model for BPD. This study also discusses the possible mechanisms involved to further understand the protective effects of UTI on lung.

## Results

### UTI treatment increased the W/D weight ratio of lung

New born rats were under hyperoxia treatment (Hyp) to induce lung injury. To quantitate pulmonary edema, the lung water content was evaluated by lung W/D weight ratio. It showed that the W/D weight ratio of Hyp group was significantly higher than Nor group. The increasing trend of W/D ratio was depressed as UTI dosage increased and the most significant decrease was observed in Hyp+UTI-H group (*p*<0.05; Fig. 1).

**Fig.1.**
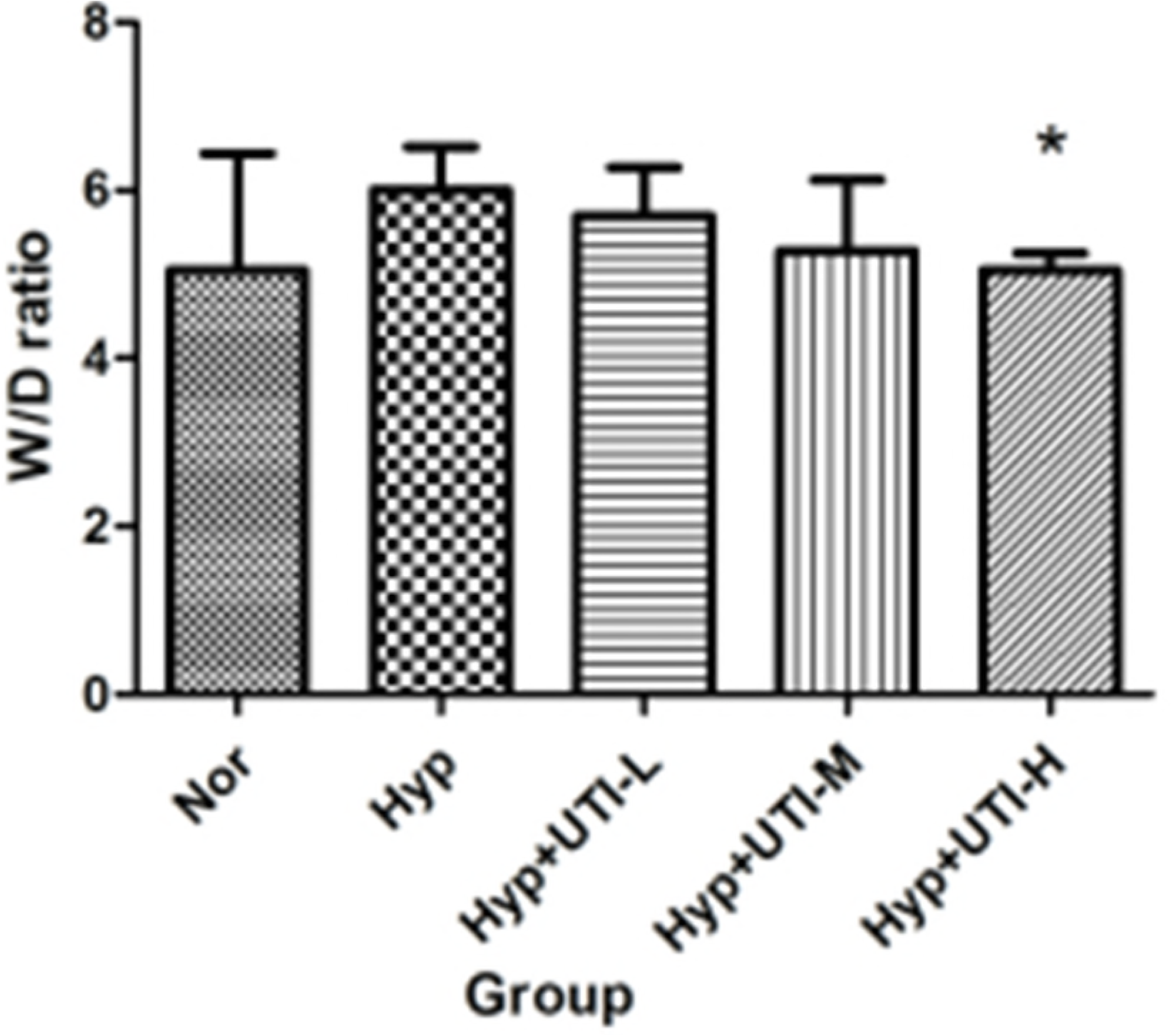
High dose of UTI treatment attenuated the W/D weight ratio of rat model of hyperoxia-induced lung injury. Histogram showing the W/D weight ratio of rats on day 41. Data are expressed in mean ± SD, \**p*< 0.05 *vs*. Hyp group.

### Effects of UTI on morphological changes of the lung induced by hyperoxia

To detect the morphological changes of lung tissues, hematoxylin and eosin staining was used and the alveolar numbers of different groups on day 41 were counted (Table 1, Fig.2). Compared to Nor group, alveoli in Hyp group rats were larger with decreased interstitial thickness and alveolar count was reduced, suggesting that hyperoxia may have inhibition effect on the formation of alveoli. The morphological changes of lung were relieved by UTI treatment in a dose-dependent fashion that the interstitial thickness, the diameter of alveolar cavity, and the number of alveoli became more similar to Nor group, especially in Hyp+UTI-H group.

**Fig.2.**
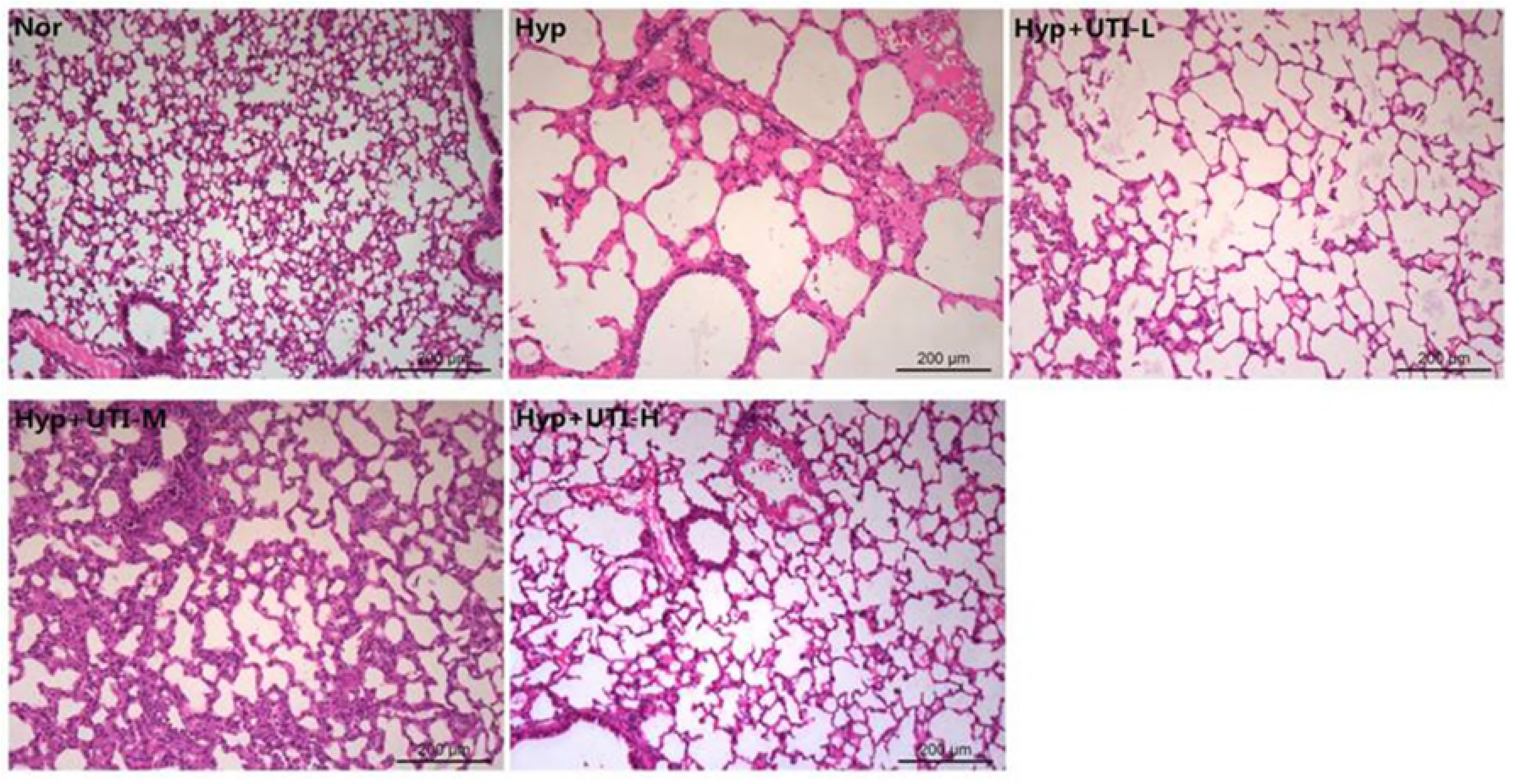
UTI treatment attenuates hyperoxia induced morphological changes of the alveoli. Pictographs showing representative pathological changes in lung tissues, from each group, as determined by hematoxylin and eosin staining. Scale bar = 200μm.

### Effects of UTI on TNF-α expression and macrophage infiltration

In order to investigate the lung inflammatory responses, the expression level of TNF-α expression was detected in different groups. As showed in Fig.3, TNF-α expression in Hyp group were higher than Nor group but UTI treatment can attenuate TNF-α level. The dramatical downregulation of TNF-α was observed in the high dose group

**Fig.3.**
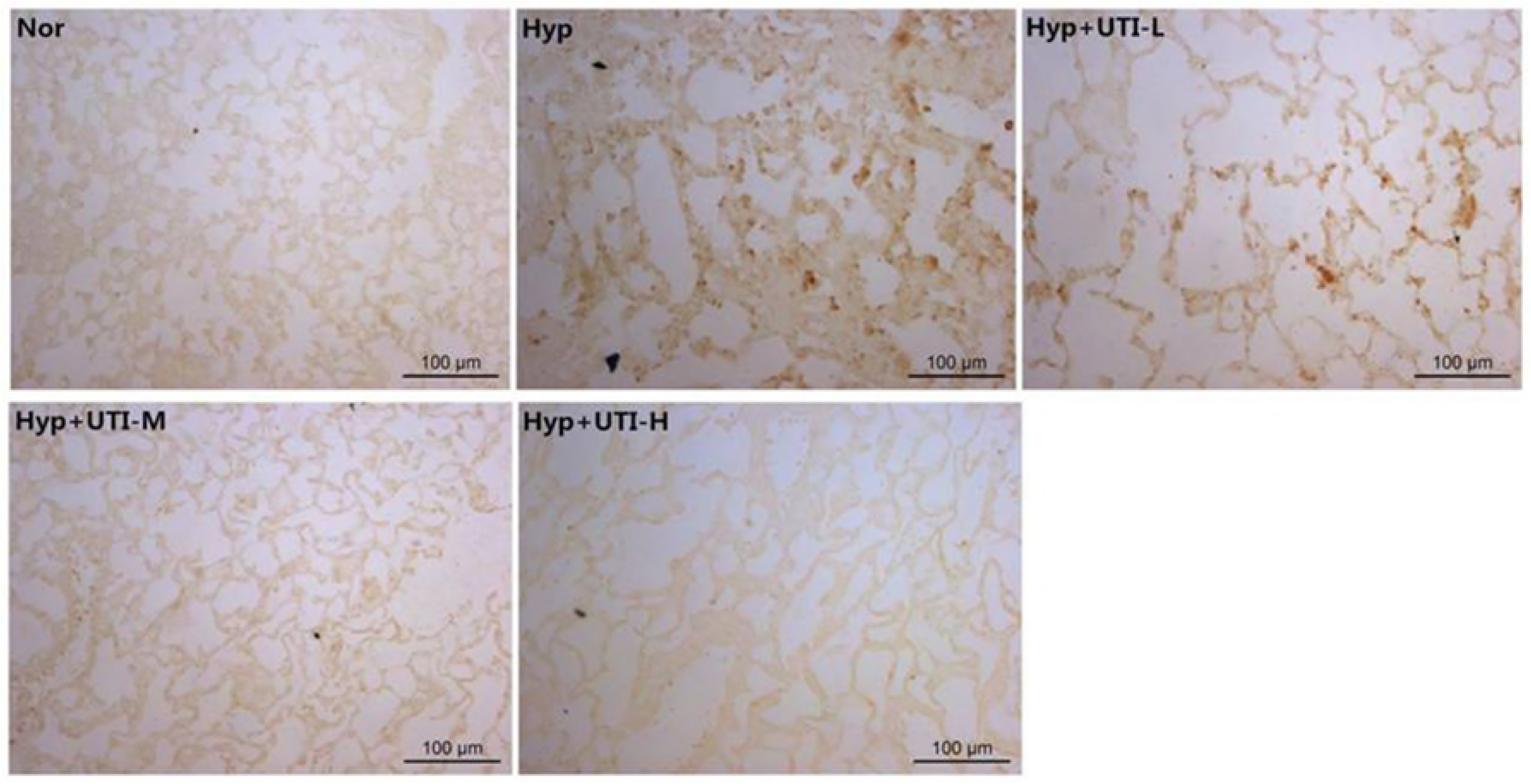
UTI attenuates hyperoxia induced TNF-α in the lungs. TNF-α expression in lung tissues were identified using immunohistochemistry, representative results are shown and scale bar = 100 μm.

We further assessed the UTI effect on macrophage infiltration through detection of CD68 expression, a marker of macrophage, by immunohistochemistry. The alveolar interstitium in Nor group was almost unstained, but it was mostly stained to dark brown in the Hyp group. The color of alveolar interstitium became moderate following UTI treatment, especially in Hyp+UTI-H group (Fig. 4). TNF-α and CD68 were mainly located around blood vessel and bronchium, and between alveolus pulmonis

**Fig. 4.**
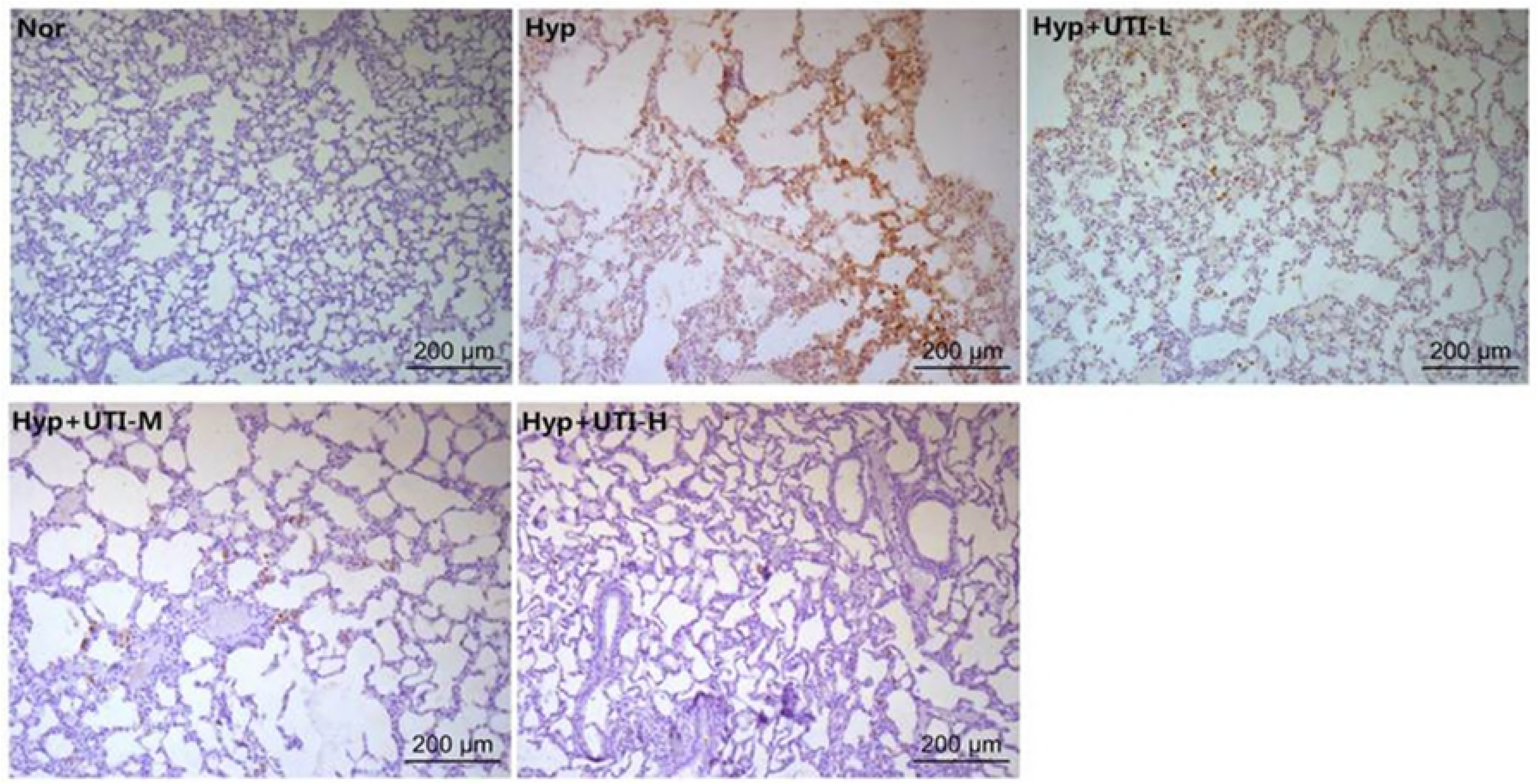
UTI inhibits hyperoxia induced macrophage infiltration of the lungs. Light micrographs showing CD68 expression in lung tissues, identified by immunohistochemistry and representative results are shown and scale bar = 200 μm.

### Effects of UTI on SOD and MDA

The effects of UTI on the oxidative stress in lung induced by hyperoxia, were detected by determination of the levels of SOD and MDA by ELISA. Compared with the Nor group, the SOD and MDA levels in lung decreased on Day 41 day after exposure to 60%±5% O_2_. However, UTI treatment had no significant effect on the levels of pulmonary SOD and MDA (Fig. 5).

**Fig. 5.**
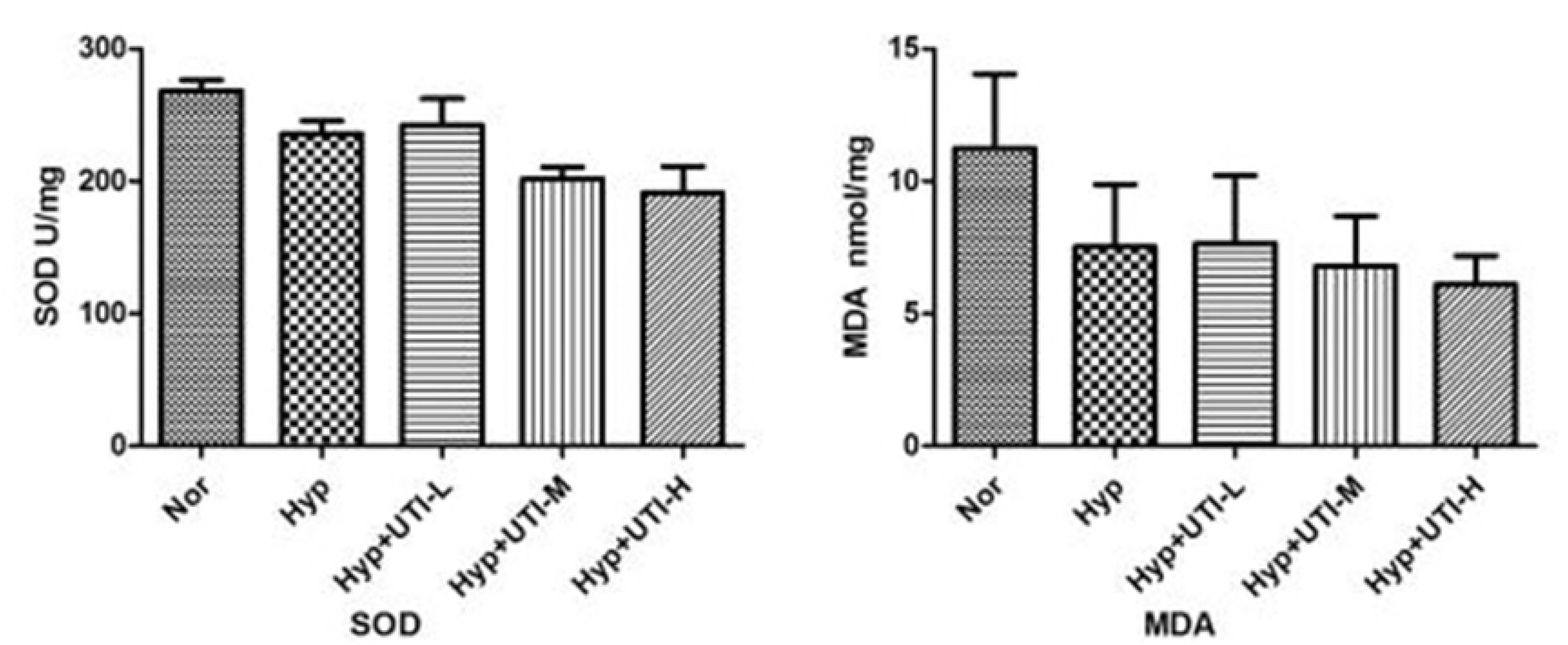
Effects of UTI on the oxidative in lung induced by hyperoxia. Histographa showing SOD and MDA levels as determined by ELISA. Data are expressed as mean ± SD.

### UTI treatment improved the average weight of rats

The average weight of rats from Hyp group was observed and compared with the control group (Nor). We found that the average weight of rats was significantly reduced in Hyp group on Day 7, and the difference was more significant on Day 21 (Table 2). Compared with the Hyp group, the average weight of rats under high dose of UTI treatment (Hyp+UTI-H) was significantly improved on Day 41, but no significant difference was observed in rats from the low dose (Hyp+UTI-L) and middle dose (Hyp+UTI-M) groups (Fig.6).

**Fig. 6.**
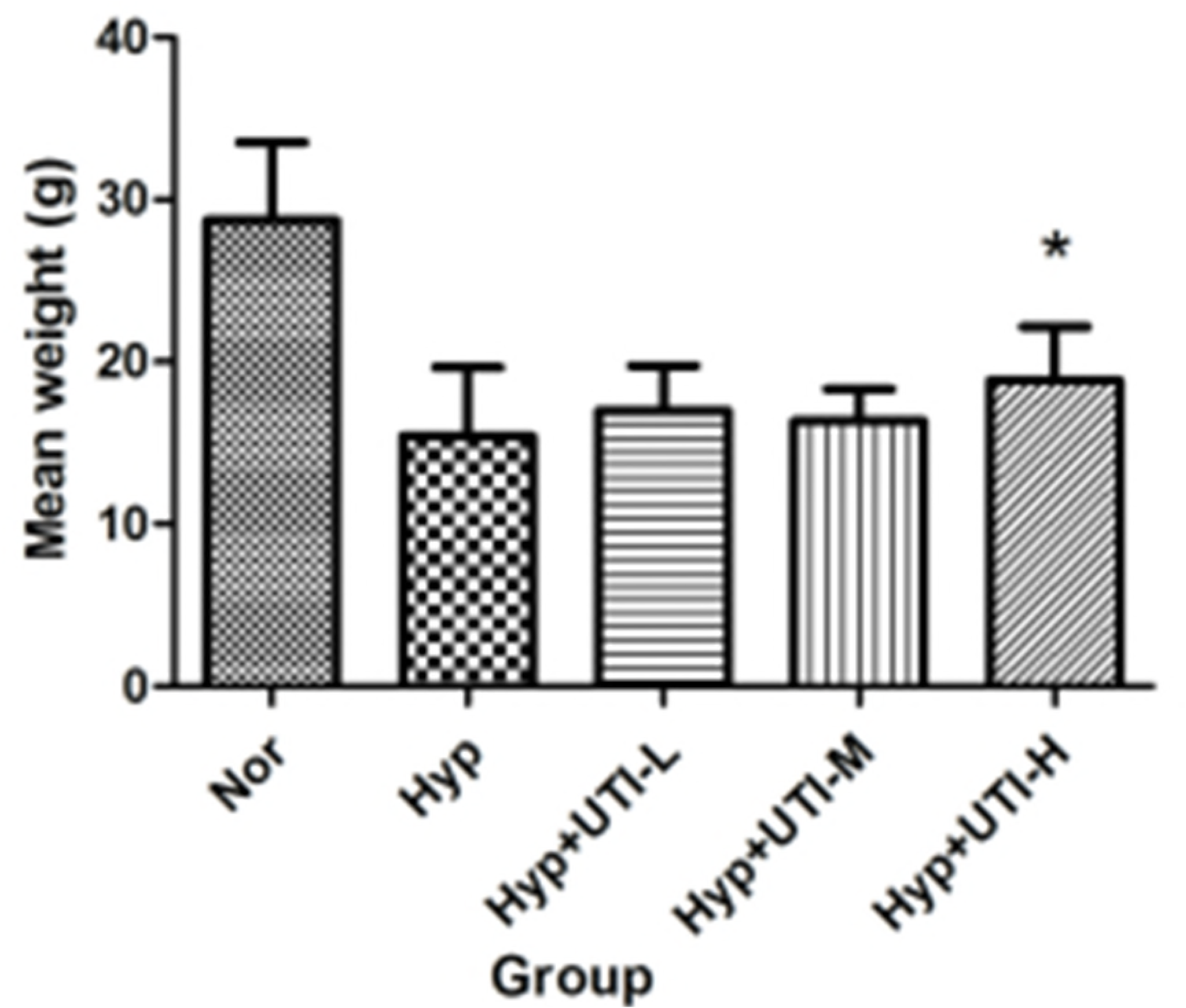
The average weights of rats of different treatments. High dose of UTI treatment improved obviously the average weight of rats of hyperoxia-induced lung injury in day 41, but no significant difference was observed in the low dose group and middle dose group. The p value was 0.048 The symbol Nor indicated rats in the normoxic control group. The symbol Hyp stood for rats treated with hyperoxia. Hyp+UTI-L meant rats in hyperoxia plus low-dose UTI treatment group. Hyp+UTI-M indicated rats in hyperoxia plus middle-dose UTI treatment group. Hyp+UTI-H indicated rats in hyperoxia plus high-dose UTI treatment group. Data were expressed in mean ± SD.

## Discussion

BPD was originally identified by Northway et al. in 1967^12^ and later redefined by Jobe ^13^ in 1999. It remains the most common complication of preterm birth which contributes significantly to morbidity and mortality but successful interventions are limited. The pathogenesis of BPD include oxidative and inflammation-mediated lung injury due to hyperoxia, ventilator-induced injury such as barotrauma and oxotrauma, infection and other stimuli. Previous studies have shown that early and sustained inflammatory cells and cytokines in tracheal fluid from premature newborns are strongly associated with high risk for the subsequent development of BPD.^14, 15^ Some studies suggest that early interventions with anti-inflammatory agents and strategies that directly preserve lung cell survival and function could prevent the development of BPD.

Hyperoxia could induce alveolar injury with endothelial cell necrosis, hemorrhage, edema, and inflammatory response as the dominant features.^16^ In this study, significant aberrations in lung development were observed in newborn rats continuously exposed to hyperoxia for 21 days, including larger alveoli, thinner interstitium, and reduced alveolar count. It has been reported that inflammatory reaction mediated by inflammatory cytokines and neutrophils has significant effects on the development of lung injury induced by hyperoxia.^2, 17^ Our results are in concert with previous studies. We observed that exposure to hyperoxia led to severe inflammatory reaction including upregulation of TNF-α, massive macrophage infiltration and pulmonary edema. It suggested that our rat models could be used in researches of hyperoxia-induced lung injury.

UTI is one of the Kunitz-type serine protease inhibitors found in human urine and blood, with heat stability and acid fastness.^18^ Previous reports have documented antioxidant and anti-inflammatory effects of UTI in animal and clinical trials. UTI inhibits intensive production of proinflammatory mediators such as IL-8, TNF-α.^19-21^ Early administration of UTI has been demonstrated to inhibit neutrophil protease release and excessive inflammatory responses and reduce the release of oxygen free radicals and consumption of superoxide dismutase.^22, 23^ Also, UTI can inhibited fibrosis formation in kidney and lungs.^24,25^ UTI has been widely used in many diseases and injuries including sepsis, severe acute pancreatitis, severe burn and multiple organ dysfunction syndrome (MODS). Thus, we hypothesized that UTI can prevent the adverse effects of hyperoxia-induced morphological changes in the lungs of newborn rats. To evaluate this hypothesis, this study explored the ability of UTI to promote pulmonary alveolar development and the mechanisms involved.

We first examined if UTI treatment had effect on pulmonary edema. In Hyp group, the lung W/D weight ratio was obviously increased and macrophage infiltration was detected. However, these changes were relieved by administration of UTI. Whether anti-inflammatory or antioxidative response played a role in protective effect against pulmonary edema of UTI was further detected.

TNF-α is largely generated by activated monocytes/macrophages and endothelial cells, which is associated with the prognosis of BPD. Molor-Erdene et al ^10^ demonstrated that UTI inhibited TNF-α and significantly reduced TNF-α expression in a dose-dependent manner in rat lung tissue. Results from present study showed that UTI treatment attenuated TNF-α level and macrophage infiltration in the lung of rats exposed to hyperoxia.

Oxidative stress is one of the important factors in lung injury induced by hyperoxia. Previous reports have documented that UTI may reduce rat lung injury and protect the lung from damage, induced by H2S, by inhibiting ROS.^11^ However, in our study, no significant differences were identified in SOD and MDA levels in Hyp groups with or without UTI treatment on Day 41. Hence, more studies are needed to further investigate the mechanisms.

We detected if UTI treatment enhanced the average body weight of rats with hyperoxia-induced lung injury and found that high dose UTI treatment significantly improved the average weight of rats, but no significant difference was observed in either the low dose group or the middle dose group.

Our study showed that UTI, in a dosage-dependent manner, protects lung of newborn rats from hyperoxia induced injury by reducing macrophage infiltration, inflammatory response, and edema. It suggested that the administration of UTI could be a potential strategy for clinical treatment of hyperoxia-induced lung injury.

## Materials and methods

### Animal model

Naturally delivered newborn Kunming rats, 2.1±0.18g in weight, were purchased from Chinese Academy of Sciences (Beijing, China). The use of animals was approved by Affiliated BaYi Children’s Hospital, Clinical Medical College in PLA Army General Hospital and conformed to the guidelines of the National Institutes of Health concerning the care and use of laboratory animals. There were 45 pups within 24 hours of birth and 5 dams randomly divided into 5 groups and each group contained 9 pups and 1 dam: hyperoxia (Hyp, 60%±5% O_2_ for 21 days + intraperitoneal saline administration for 21 days); hyperoxia plus low-dose UTI (Hyp+UTI-L, 60%±5% O_2_ for 21 days + intraperitoneal UTI 10,000U/kg administration for 21 days); hyperoxia plus middle-dose UTI (Hyp+UTI-M, 60%±5% O_2_ for 21 days + intraperitoneal UTI 50,000U/kg administration for 21 days); hyperoxia plus high-dose UTI (Hyp+UTI-H, 60%±5% O_2_ for 21 days + intraperitoneal UTI 100,000U/kg administration for 21 days); Normoxic control (Nor, 21% O_2_ for 21 days+ intraperitoneal saline administration for 21 days). Hyperoxia treatment was conducted as previously described^9^ that the rats were continuously supplied with oxygen in oxygen chambers (FiO_2_ = 60%±5%, monitored by oxygen recorder) and with constant temperature (22°C∼26°C) and humidity (50%-60%). The dosage and schedule of UTI used in the following publications were referred to our study.^10,11^ Every day, the chambers were opened at fixed time to perform water and food supply and drug treatment, and all these operations were conducted within 1 hour. Maternal rats were daily exchanged with rat from control group to avoid poor feeding influenced by oxygen toxicity. 10,000 U/kg, 50,000 U/kg and 100,000 U/kg of UTI was injected intraperitoneally every day to hyperoxia plus low-dose UTI, hyperoxia plus middle-dose UTI and hyperoxia plus high-dose UTI groups, respectively, from the 21th day after birth. Pups in hyperoxia group were administered the same volume of solvent from the 21^st^ day after birth. Pups in normoxic control group were exposed to normoxia (FiO_2_ = 21%) and administered the same volume of solvent from the 21^st^ day after birth, once daily. Weights of the rats were recorded every other day. And after the 41-day experimental period, the animals were sacrificed to harvest lung samples.

### Lung wet/dry (W/D) weight ratio

The upper part of the left lung lobe was weighed (wet weight, W) and then dried to constant weight (dry weight, D) in a thermostatic oven at 60°C for 72h. The tissue edema and the W/D weight ratio were subsequently calculated.

### Histopathological examination

The lower part of the left lung lobe was embedded with paraffin using conventional procedures, then sliced into 5-μm thickness. Pathological studies were conducted through hematoxylin and eosin staining. For immunohistochemical analysis, the sliced lung tissue was rinsed with 1% phosphate buffered saline. The slides were treated with 0.3-3% H_2_O_2_ in methanol for 12 min and washed with PBS for three times. Antigen retrieval was performed at 92°C for 10 min. After washing with PBS, the slides were treated with goat serum then incubated with primary antibody at 37°C overnight, second antibody at 37°C for 40 min, and HRP-conjugated anti-mouse IgG at 37°C for 30 min. DAB and hematoxylin staining was respectively performed for 10 min and 30 sec, which was terminated with ddH_2_O. Then the slides were washed, dehydrated, and coverslipped. Rabbit anti-TNF-α and rabbit anti-CD68 antibodies (Abcam) were 1:1000 diluted.

### ELISA analysis

The right lobes of the lung tissues were immediately frozen in liquid nitrogen and stored at −80°C. The protein levels of superoxide dismutase (SOD) and malondialdehyde (MDA) were assessed using corresponding ELISA kit (Boster). The total protein from rat lung tissue was extracted to measure protein content using the BCA (bicinchoninic acid) method.

### Statistical analysis

The SPSS version 19.0 statistical software was used for data analyses. All data are presented as mean ± standard deviation (SD). Data from different groups were compared using one-way analysis of variance (ANOVA) followed by Scheffé multiple comparison test. The Student’s t test was used for pairwise comparisons. When *p was* <0.05, the difference between different groups was considered significant.

## Acknowledgements

None.

